# Ethylene signaling induces gelatinous layers with typical features of tension wood in hybrid aspen

**DOI:** 10.1101/204602

**Authors:** Judith Felten, Jorma Vahala, Jonathan Love, András Gorzsás, Markus Rüggeberg, Nicolas Delhomme, Joanna Leśniewska, Jaakko Kangasjärvi, Torgeir R. Hvidsten, Ewa J. Mellerowicz, Björn Sundberg

**Author notes:** **Corresponding author details:** Judith Felten.

## Abstract

**Research conducted:** The phytohormone ethylene impacts secondary stem growth in plants by stimulating cambial activity, xylem development and fiber over vessel formation. Here we report the effect of ethylene on secondary cell wall formation and the molecular connection between ethylene signaling and wood formation.

**Methods:** We applied exogenous ethylene or its precursor 1-aminocyclopropane-1-carboxylic acid (ACC) to wild type and ethylene insensitive hybrid aspen trees *(Populus tremula x tremuloides)* and studied secondary cell wall anatomy, chemistry and ultrastructure. We furthermore analyzed the transcriptome (RNA Seq) after ACC application to wild type and ethylene insensitive trees.

**Key results:** We demonstrate that ACC and ethylene induce gelatinous-layers (G-layers) and alter the fiber cell wall cellulose microfibril angle. G-layers are tertiary wall layers rich in cellulose, typically found in tension wood of aspen trees. A vast majority of transcripts affected by ACC are downstream of ethylene perception and include a large number of transcription factors (TFs). Motif-analyses reveal potential connections between ethylene TFs (ERFs, EIN3/EIL1) and wood formation.

**Conclusion:** G-layer formation upon ethylene application suggests that the increase in ethylene biosynthesis observed during tension wood formation is important for its formation. Ethylene-regulated TFs of the ERF and EIN3/EIL1 type could transmit the ethylene signal.

## Introduction

Tension wood (TW) is formed on one side of a stem or a branch of angiosperm trees upon displacement resulting from, for example, bending or leaning. TW induces tensional stress in the wood that reorients the branch/stem towards its original position (Mellerowicz & Gorshkova, 2012; Felten & Sundberg, 2013; Fagerstedt *et al.*, 2014; Fournier *et al.*, 2014; Ruelle, 2014; Groover, 2016). TW is associated with an asymmetric cambial growth of the stem/branch, decreased size and frequency of vessels, and a change in the ultrastructure and chemistry of the fiber cell wall. A particular feature of TW in many angiosperm trees is the occurrence of gelatinous fibers (G-fibers), characterized by a tertiary gelatinous cell wall (G-layer) on the inner (lumen) side of the fiber secondary cell wall (SCW) layers. The G-layer is highly enriched in cellulose, has insignificant lignin content, a few matrix polysaccharides that are distinctively different from the polysaccharides of adjacent S-layers, and a much steeper cellulose microfibril angle (MFA) compared to that in the adjacent S2 layer of the G-fiber (Goswami *et al.*, 2008; Clair *et al.*, 2011; Felten & Sundberg, 2013).

Ethylene is thought to be an upstream trigger of the TW response: ethylene biosynthesis increases during TW formation, and applied ethylene induces some of the typical TW attributes, such as increased cambial growth and decreased vessel size and frequency (Andersson-Gunnerås *et al.*, 2003; Du & Yamamoto, 2007; Love *et al.*, 2009). The asymmetric growth response in leaning stems requires functional ethylene signaling and is absent in ethylene-insensitive hybrid aspen trees (Love *et al.*, 2009). However, conclusive information on how ethylene signaling impacts the properties of the fiber cell wall and, in particular, induction of G-layers is lacking.

The effects of ethylene on wood formation imply that its downstream signaling affects important regulators of wood development. Ethylene is perceived by ethylene receptors (ETRs) at the endoplasmic reticulum, which trigger downstream signaling pathways involving phosphorylation, ubiquitination and translocation, hence activating a set of transcription factors (TFs), namely ETHYLENE INSENSITIVE 3/ETHYLENE INSENSITIVE3-LIKE1 (EIN3/EIL1) and Ethylene Response Factor (ERF) TFs. These TFs control their respective targets in diverse ethylene response pathways (Alonso & Stepanova, 2004; Merchante *et al.*, 2013). ERF proteins regulate targets in various developmental processes by binding to GCC boxes in the promoter of their targets, which usually enhances their transcription (Ohmetakagi & Shinshi, 1995; Shinshi *et al.*, 1995; Büttner & Singh, 1997; Allen *et al.*, 1998). EIN3/EIL TFs are known to bind to the TEIL motif in the promoters of ERFs and diverse immediate downstream targets, and to regulate their expression (Solano *et al.*, 1998; Kosugi & Ohashi, 2000). Therefore both ERFs and EIN3/TEIL TFs represent a link connecting ethylene signaling to wood development. Indeed, in Arabidopsis AtERF1, ERF018 and ERF109 are required for the increase in cambial cell division observed in ethylene overproducing *eto* mutants, providing evidence that ERFs are important for ethylene signaling in vascular tissues (Etchells *et al.*, 2012). Moreover, overexpression of certain *Populus* ERFs in transgenic hybrid aspen leads to an overall modification of the carbohydrate to lignin ratio in wood (Vahala *et al.*, 2013). However, the detailed molecular connection between ERFs, or general ethylene effects, and wood formation remains elusive.

Here we study ethylene effects on fiber cell walls and transcriptional signaling downstream of ethylene perception by using ethylene insensitive hybrid aspen and two experimental systems established by Love et al. (2009). In the first system, ethylene production was stimulated by supplying the ethylene precursor 1-aminocyclo-propane-1-carboxylate (ACC) to the root system of *in vitro*-grown hybrid aspen. Thereby, ACC reached the woody tissues through the xylem transpiration stream. In the second system, an ethylene flow-through cuvette was mounted around one internode per greenhouse-grown tree to apply ethylene gas. The use of ethylene insensitive hybrid aspen trees allowed us to confirm that the observed ACC and ethylene effects on wood formation and the transcriptome responses to ACC require ethylene signaling.

## Material and Methods

### Plant Material, in vitro growth conditions and ACC treatment

Plant material was wild type hybrid aspen (*Populus tremula* L. x *tremuloides* Michx; clone T89) or transgenic ethylene-insensitive hybrid aspen lines *p35S::etr1-1* (line 1E) and *pLMX5::etr1-1* (line 6) previously described (Love *et al.*, 2009) and treated with 100 μM ACC in a procedure similar to the one described in Love et al. (2009), for details see Method S1. For sampling, the fourth internode below the apex (reference internode (RI)), from which secondary growth starts (Love et al. 2009), was marked at the beginning of the ACC treatment. For RNA extraction and transcript analysis, entire stems from six plants per treatment and genotype were harvested 10 h after ACC or water application from the bottom of the stem up to the RI and frozen in liquid N_2_. For all other analyses, plants were allowed to grow for two weeks in three replicate experiments (Set1, Set2, Set3) from the starting point of treatment under the above-mentioned growth conditions. Histological analysis was carried out for each set with at least five biological replicates per treatment and genotype. All other analyses were carried out with plants from two sets (Set1 and Set3) with all material for one type of analysis originating always from plants of the same set. Material was collected close to the RI to ensure the highest possible proportion of wood formed during the treatment.

### Ethylene treatments and material sampling

The two-week ethylene treatment was carried out by applying synthetic air (with 380 ppm of carbon dioxide) with ethylene (2 ppm) or without (control) through flow-through cuvettes on the stems of greenhouse-grown hybrid aspen trees (three-months old and approximately 2 m tall) as described in Love et al. (2009). Three replicate trees for each genotype (wild type, *p35S::etr1-1* and *pLMX5::etr1-1*) were subjected to the respective treatment. At the end of the treatment the ethylene-treated internodes with respective controls were harvested, cut into three parts (bottom, middle, top) and stems were halved longitudinally. Material for all three internodes was used for embedding and light or transmission electron microscopy. Because no major differences depending on sampling position throughout the exposed internode were observed (Fig. S1), only material from the middle part of the treated internode was analyzed by microscopy and X-ray diffraction.

### Histological analysis

Histological investigation of *in vitro*-grown trees was carried out on transverse fresh hand-cut sections stained with safranin/alcian blue. Fixed, embedded (see below) and ultramicrotome-cut 1 μm thick sections from control/ethylene-treated stems of greenhouse-grown trees were stained with toluidine blue. All sections were examined using a Zeiss Axioplan 2 microscope. Immunolocalization was performed as outlined in Method S2.

### Transmission electron microscopy

Approximately 0.5 mm thick transverse hand-cut sections from *in vitro*- or greenhouse-grown material (3^rd^ internode below RI from Set1 for ACC treatments; bottom, middle and top part of exposed internodes for ethylene gas treatments) were fixed in 2.5% glutaraldehyde in sodium cacodylate buffer (0.1M, pH7.2) for > 48 h, treated for 2 h with 1% OsO_4_ and then processed and imaged as described in (Roach *et al.*, 2012).

### Microfibril angle measurements by X-ray diffraction

The third internode below RI (Set3) of water- and ACC-treated plants and the middle segment of ethylene-treated greenhouse-grown plants with respective controls were frozen in liquid N_2_. Radial-longitudinal sections of 100 μm thickness were cut with a cryotome and dried between two microscopy glass slides. Xylem strips were transferred to metal holders for X-ray diffraction analysis. The X-ray diffraction experiments were carried out with two different Bruker Nanostar (Bruker AXS, Karlsruhe, Germany) devices, both equipped with a Hi-Star area detector and using Cu Kα radiation (8 keV, λ=0.154 nm). The beam diameter of one device was set approximately to 150 μm, whereas that of the second device was set to approximately 400 μm. The (200)-Bragg peaks of cellulose, which were taken for cellulose orientation analysis, appear at a scattering angle 2 *θ* of 22.6°. The samples were measured in an evacuated chamber with the long axis of the fiber cells being perpendicular to the incident X-ray beam. The sample-detector distance was set to 68 mm. For the *in vitro*-grown plants and for most of the greenhouse-grown plants the device with the 400 μm beam was used, whereas for the ethylene-treated wild type greenhouse-grown plants, the 150 μm diameter beam was used to achieve a higher spatial resolution of the measured area along the radial axis of the sample. The sample width was between 0.5 and 2 mm, so several measurements could be performed on one sample with an exposure time of 1 h. Azimuthal intensity profiles (azimuthal angle φ vs. intensity) of the diffraction patterns were obtained by radially integrating the intensity of the (200)-peaks within 2θ±0.2° with an azimuthal step size of 1°. Microfibril orientation distributions and mean MFA were calculated by the simulation and fitting routine as described in detail previously (Rüggeberg *et al.*, 2013).

### Fourier-transform infrared (FT-IR) microspectroscopy

FT-IR microspectroscopy was performed using existing protocols (Gorzsás *et al.*, 2011; Gorzsás & Sundberg, 2014), for technical details see Method S3. Eighteen micrometer thick transverse sections were prepared from frozen samples (fourth internode below the RI, Set3) using a Microm cryotome HM505E, and dried in a desiccator between two microscopy glass slides for at least 48 h. Two sections from each of four biological replicates per genotype and treatment were analyzed. Hyperspectral images were recorded over two (ACC-treated plants) or one (water-treated plants) position(s). Five spectra per image were extracted from selected positions, corresponding to the blue ring in ACC samples and a similar position in water treated controls. Multivariate analyses (principal component analysis, PCA, and orthogonal projections to latent structures discriminant analysis, OPLS-DA (Trygg & Wold, 2002)) were performed using SIMCA-P+ (version 12.0, Umetrics AB, Umeå, Sweden), for explanation see Method S3.

### Transcriptomics by RNA sequencing

RNA was extracted from the stems of wild type, *p35S::etr1-1* and *pLMX5::etr1-1* hybrid aspen trees treated with ACC or water for 10 h, with three replicates per condition and genotype. Each sample (replicate) was a pool of two plants. Material from entire stems below RI was ground with mortar and pestle in liquid N_2_. One hundred mg of frozen stem powder was used for CTAB RNA extraction followed by lithium chloride precipitation (Chang *et al.*, 1993). RNA was resuspended in 50 μl DEPC-treated milliQ water. One part of the samples was diluted to 250 ng/μl in 50 μl and residual DNA was removed using *DNAfree*^™^ (Ambion) according to the manufacturer’s instructions. After DNA removal, RNA was cleaned using the Qiagen MinElute kit. RNA concentration was measured with a Nanodrop ND-1000 (Nano-Drop Technologies, DELAWARE, USA) and its quality was assessed using an Agilent 2100 Bioanalyzer with Agilent RNA 6000 Nano Chips according to the manufacturer’s instructions.

Sequencing library generation and sequencing using Illumina HiSeq 2000 were carried out at SciLifeLab (Science for Life Laboratory, Stockholm, Sweden). Further information about the sequencing and data analysis procedure is available in Method S4. An overview of the data, including raw and post-QC read counts and alignment rates, is given in Table S1. Raw sequencing data are available at the European Nucleotide Archive (ENA) https://www.ebi.ac.uk/ena under the accession number ERP012528.

Generation of Venn Diagrams for differentially expressed genes in all three genotypes (wild type and ethylene-insensitive hybrid aspen) was performed in R (VennDiagram package). Gene Ontology (GO) enrichment analysis was performed with the GO enrichment tool at www.popgenie.org (Sundell *et al.*, 2015).

### Motif analysis

Promoter regions of lengths 500 bp, 1 kbp, 1.5 kbp and 2 kbp were extracted upstream of the start codon of genes in the *P. trichocarpa* genome (39,256 genes had extractable promoters). Exact matches of the EIN3 targeted TEIL motif (AYGWAYCT where Y is A or C and W is A or T (Kosugi & Ohashi, 2000)) and the ERF targeted GCC box (AGCCGCC) (Ohmetakagi & Shinshi, 1995; Shinshi *et al.*, 1995; Büttner & Singh, 1997; Allen *et al.*, 1998) within these promoter regions were identified on both strands. Statistically significant enrichment of motif matches in different subsets of differentially expressed genes was computed using the Hypergeometric distribution.

## Results

### Ethylene signaling induces formation of gelatinous cell wall layers in xylem fibers

Application of ACC to the medium of *in vitro*-grown hybrid aspen plants stimulated cambial growth and resulted in a xylem with smaller and fewer vessels. This response was absent in the ethylene-insensitive lines *pLMX5::etr1-1* and *p35S::etr1-1* (Fig. 1 and Fig. S4), in line with previous results (Love *et al.*, 2009). The ACC treatment also modified the xylem cell wall chemistry. Safranine/alcian blue, which stains matrix polysaccharides in blue and lignin in red, marked a blue circumferential band in the central part of the xylem (Fig. 1b and Fig. S4). This band was induced by ACC in the wild type but not observed in either of the ACC-treated ethylene-insensitive lines, indicating its dependence on ethylene signaling. In a few samples of ACC-treated ethylene-insensitive trees, however, small disconnected patches of blue stained fibers could be observed, which we attribute to a slight leakiness in ethylene insensitivity. Interestingly, in ACC-treated wild type trees, the blue band was only produced during an initial phase after the ACC treatment, whereas the ACC-induced decrease in vessel frequency was maintained during the two-week experimental period (Fig. 1).

**Figure 1.**
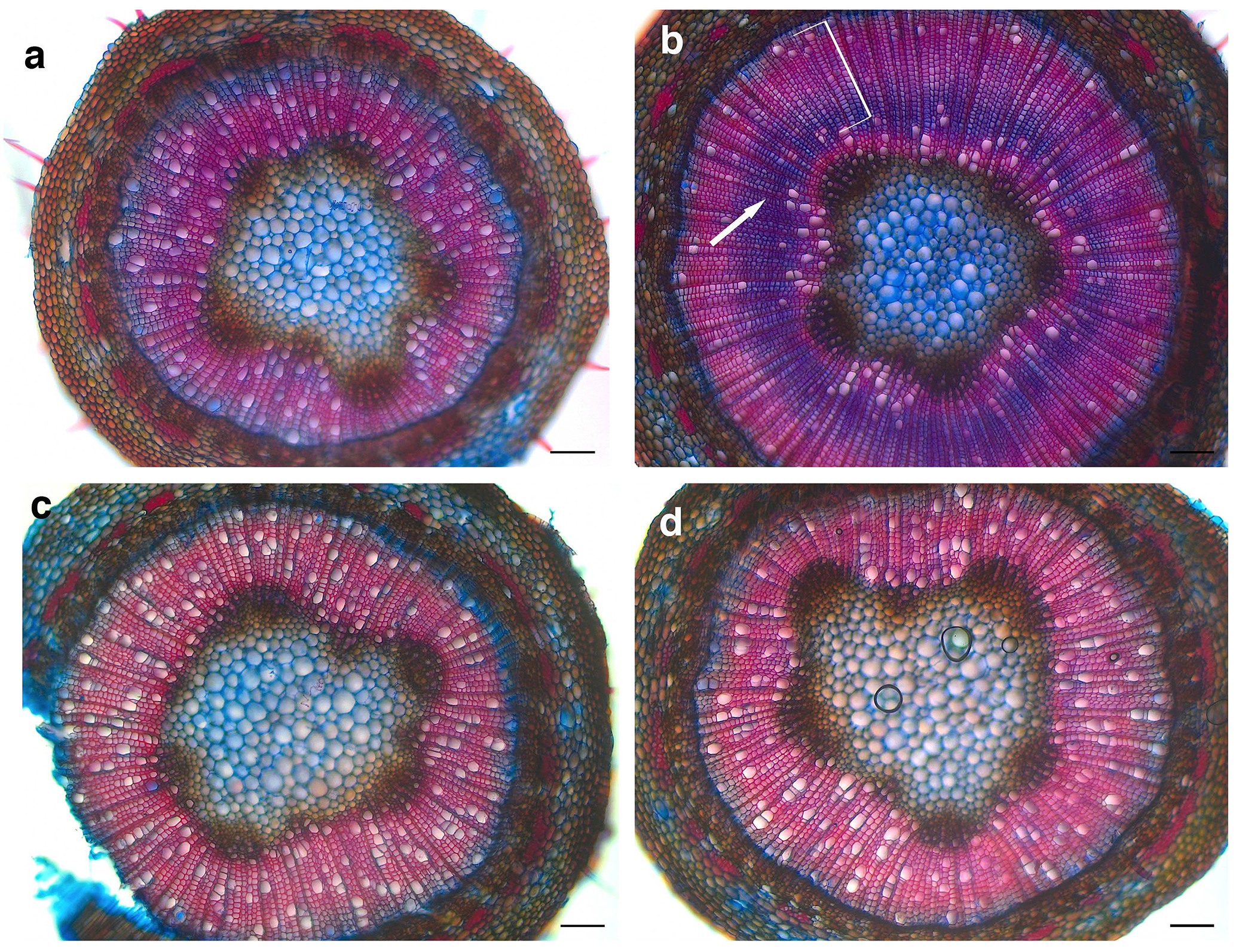
Light microscopy images of safranin/alcian blue-stained cross-sections of *P. tremula x P. alba* showing (a) a water-treated wild type tree, (b) an ACC-treated wild type tree, (d and d) an ethylene-insensitive tree (*p35S::etr1-1* and *pLMX5::etr1-1*, respectively). Note the blue circumferential staining (arrow) and a region with reduced size and frequency of vessels (bracket) induced by ACC treatment (in b) that are absent in the ethylene-insensitive trees (c and d). Scale bars =100 μm.

At higher magnification, it could be seen that the blue stain induced by ACC was observed at the lumen side of the S-layer (Fig. 2), whereas the surrounding S-layers exhibited red (lignin) staining. Transmission electron microscopy analysis confirmed the existence of the additional cell-wall layer with typical G-layer morphology, easily detached from the S-layers (Fig. 2b-d and Fig. S5). Moreover, immunolocalization of β-1,4 galactan and xylan with the monoclonal antibodies LM5 and LM10 showed that the fibers located in the blue xylem band had a high β-1,4,-galactan content, which is specific to G-fibers, whereas the fibers at the periphery of the blue band had a high xylan content that is typical for S-fibers in aspen (Mellerowicz & Gorshkova, 2012; Gorshkova *et al.*, 2015) (Fig. S6). Thus, the blue band of fibers formed after ACC treatment shows features typical of G-fibers in TW (Norberg & Meier, 1966; Clair *et al.*, 2005; Chang *et al.*, 2009; Felten & Sundberg, 2013).

**Figure 2.**
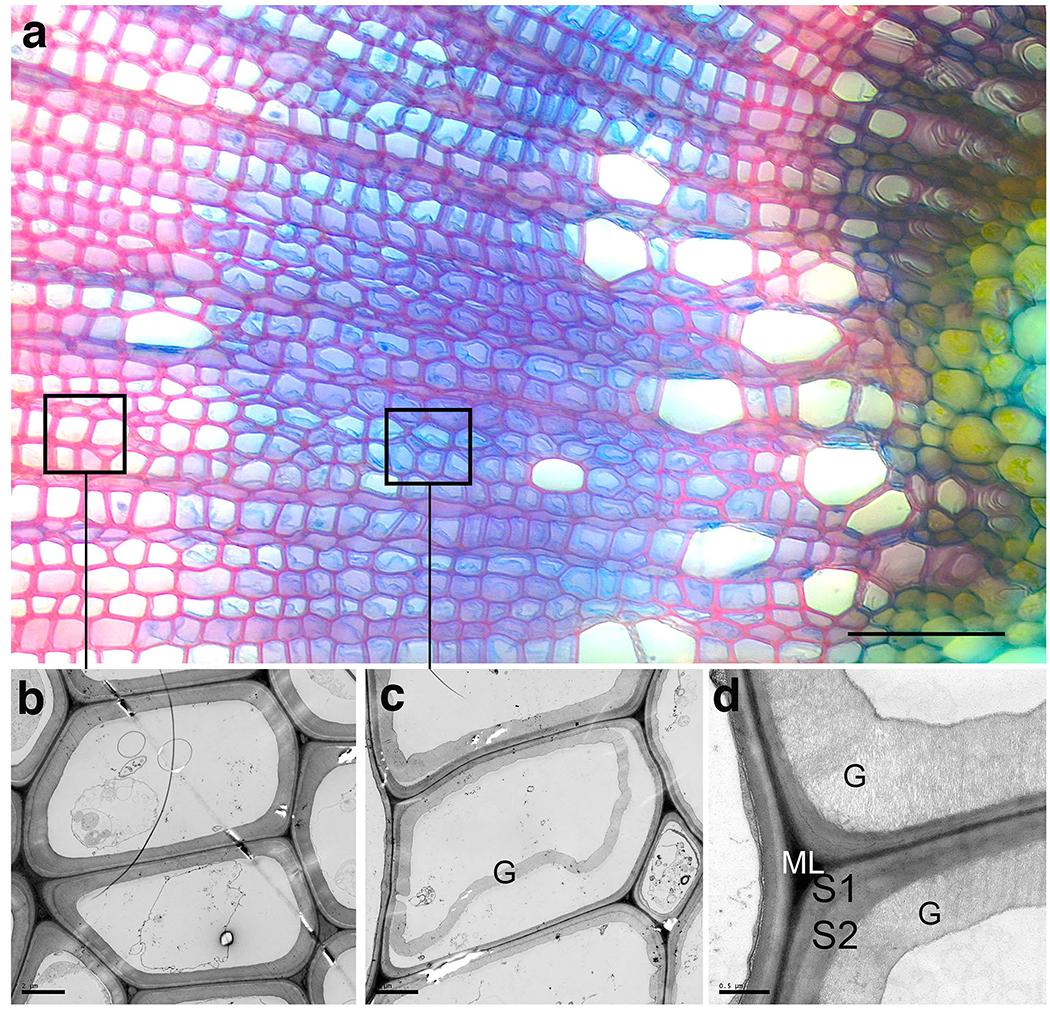
(a) Light microscopy image of a safranin/alcian blue -stained cross-section of *P. tremula x P. alba* showing fibers with G-layers (blue) in wild type tree treated with ACC. (b) TEM analysis of fibers at the xylem-periphery showing the absence of G-layers in the xylem, (c) fibers towards the center of the stem show the presence of G-layers (denoted G) partially detached from the S2-layer, and (d) magnification of a cell corner from G-fibers, where the porous structure of the G-layer (G) can be seen; middle lamella (ML) and S1-and S2-layers are separated. Scale bars = 50 μm (a), 2 μm (b, c) and 0.5 μm (d).

OPLS-DA models based on Fourier-transform infrared (FT-IR) microspectroscopy data (according to Gorzsás et al. (2011)) support the ACC effect on the cell wall chemistry specifically in the G-fiber band of ACC-treated wild type plants (Fig. 3a). Such effects were absent in the ethylene-insensitive line *pLMX5::etr1-1*, and strongly decreased in line *p35S::etr1-1* (Fig. 3c, e), consistent with earlier findings that in woody tissues *pLMX5::etr1-1* exhibits greater insensitivity to ACC than *p35S::etr1-1* (Love *et al.*, 2009). ACC treatments of wild type trees induced a proportional increase in bands indicative of carbohydrates and a proportional decrease in bands indicative of aromatics (e.g. lignin), in line with the presence of a G-layer (Fig. 3b). The fact that small patches of G-fibers were also occasionally observed in ACC-treated ethylene-insensitive plants explains why similar Loadings plots are obtained for ACC-treated *p35S::etr1-1* and *pLMX5::etr1-1* as for the wild type (Fig. 3d, f), even though Scores Plots indicated that the chemical differences between ACC and water treatment in these lines is much weaker than in wild type plants.

**Figure 3.**
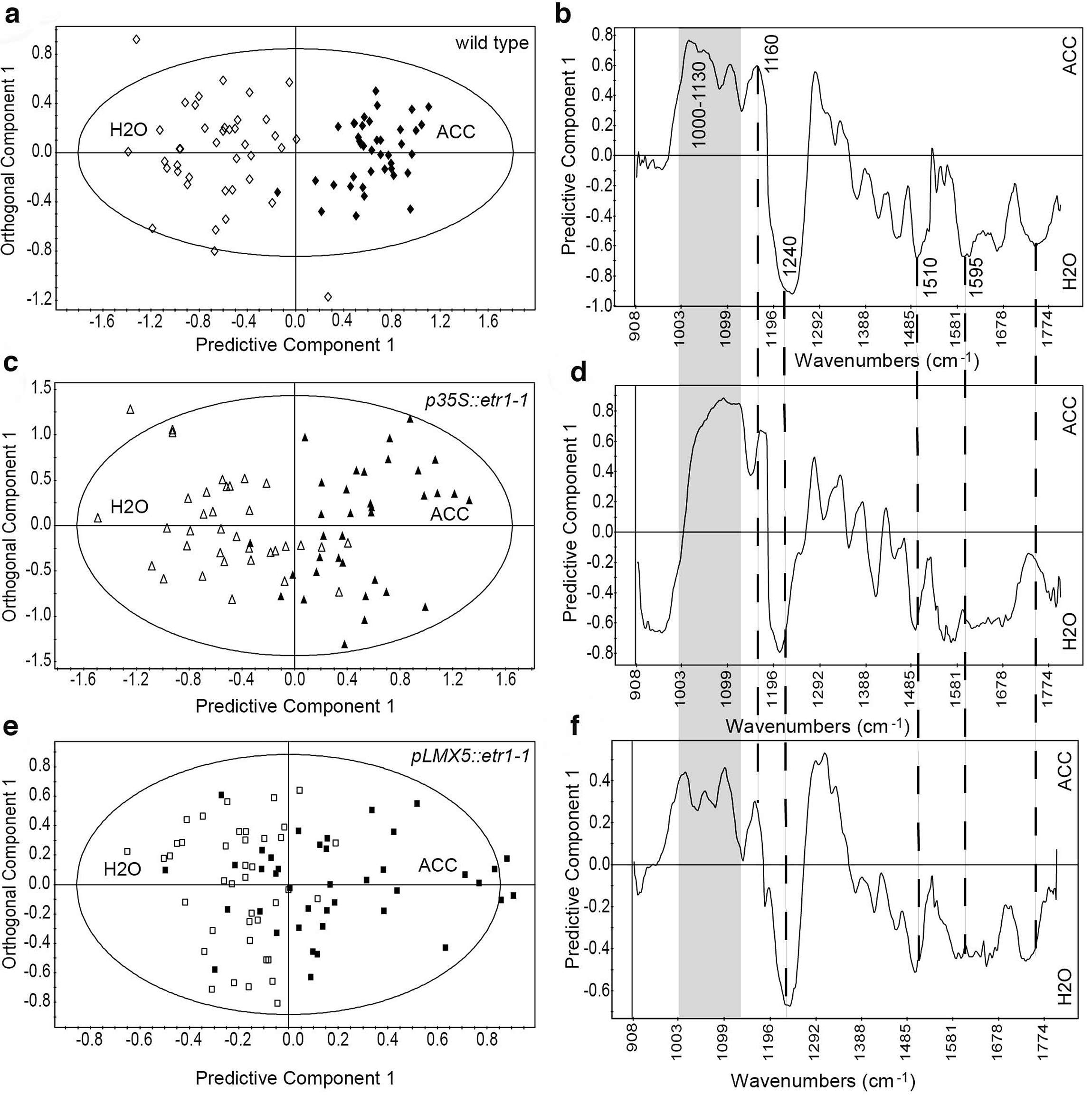
Chemical analysis by FT-IR microspectroscopy of gelatinous fibers in ACC-treated *P. tremula x P. alba* wild type trees, and of fibers in corresponding positions in water (H_2_O)-treated wild type trees, and ACC-and water-treated ethylene-insensitive trees. OPLS-DA Scores plots in (a) wild type, (c) *p35S::etr1-1* and (e) *pLMX5:etr1-1* plants, showing the separation between water-treated control (empty symbols) and ACC-treated (filled symbols) fibers. Each symbol represents spectra obtained from one fiber. Spectra were collected from four replicate plants per genotype and treatment, with ten spectra (originating from two different positions in each section) per treatment and plant. The corresponding Loadings plots for the predictive components, showing factors separating ACC-treated fibers from water-treated controls in (b) wild type, (d) *p35S::etr1-1* and (f) *pLMX5:etr1-1* plants. Bands with positive Loadings are more intense in ACC-treated fibers, whereas bands with negative Loadings are more intense in water-treated controls. The marked bands represent the following: 1000-1130 cm^−1^: unspecific carbohydrate vibrations; 1160 cm^−1^: asymmetric-C-O-C-stretch; 1240 cm^−1^:-C-O vibration; 1510 and 1595 cm^−1^: aromatic-C=C-vibrations; 1740 cm^−1^:-C=O vibration. All models have one predictive and two orthogonal components. Model details are: wild type (A, B): R2X(cum) = 0.832, R2Y(cum) = 0.805, Q2(cum) = 0.765; *p35S::etr1-1* (c, d) R2X(cum) = 0.847, R2Y(cum) = 0.621, Q2(cum) = 0.542; *pLMX5::etr1-1* (e, f), R2X(cum) = 0.816, R2Y(cum) = 0.335, Q2(cum) = 0.182.

The ability of ACC to induce G-fibers with a typical cellulose microfibril orientation distribution was examined by X-ray diffraction according to Rüggeberg et al. (2013). The S2 cell wall layer in juvenile wood in upright-grown hybrid aspen stems has a typical MFA of about 5-15° (Bjurhager *et al.*, 2010; Svedström *et al.*, 2012), whereas G-layers of TW have very low MFA of about 0-5° (Clair *et al.*, 2011). Moreover, a bimodal distribution of the cellulose microfibrils is observed in TW corresponding to the G-layer and the S2-layer, with the latter having an MFA between 20 and 40° in TW (Müller *et al.*, 2006; Goswami *et al.*, 2008). The xylem from *in vitro*-grown plants was measured with a 400 μm diameter beam that spanned the entire xylem area and therefore, the obtained data contain, in the case of ACC-treated wild type plants, contributions from areas both with and without G-layers. The MFA of the ACC-treated wild type plants exhibited a bimodal MFA distribution pattern with maxima at 3° and 29°, indicative of typical G-fibers (Fig. 4). In contrast, the MFA distribution was uni-modal in all other samples, indicating absence of G-layers, and yielded MFAs (as maxima of the distribution) in the range of 4° to 9° for water-treated wild type plants and water- or ACC-treated ethylene-insensitive lines.

**Figure 4.**
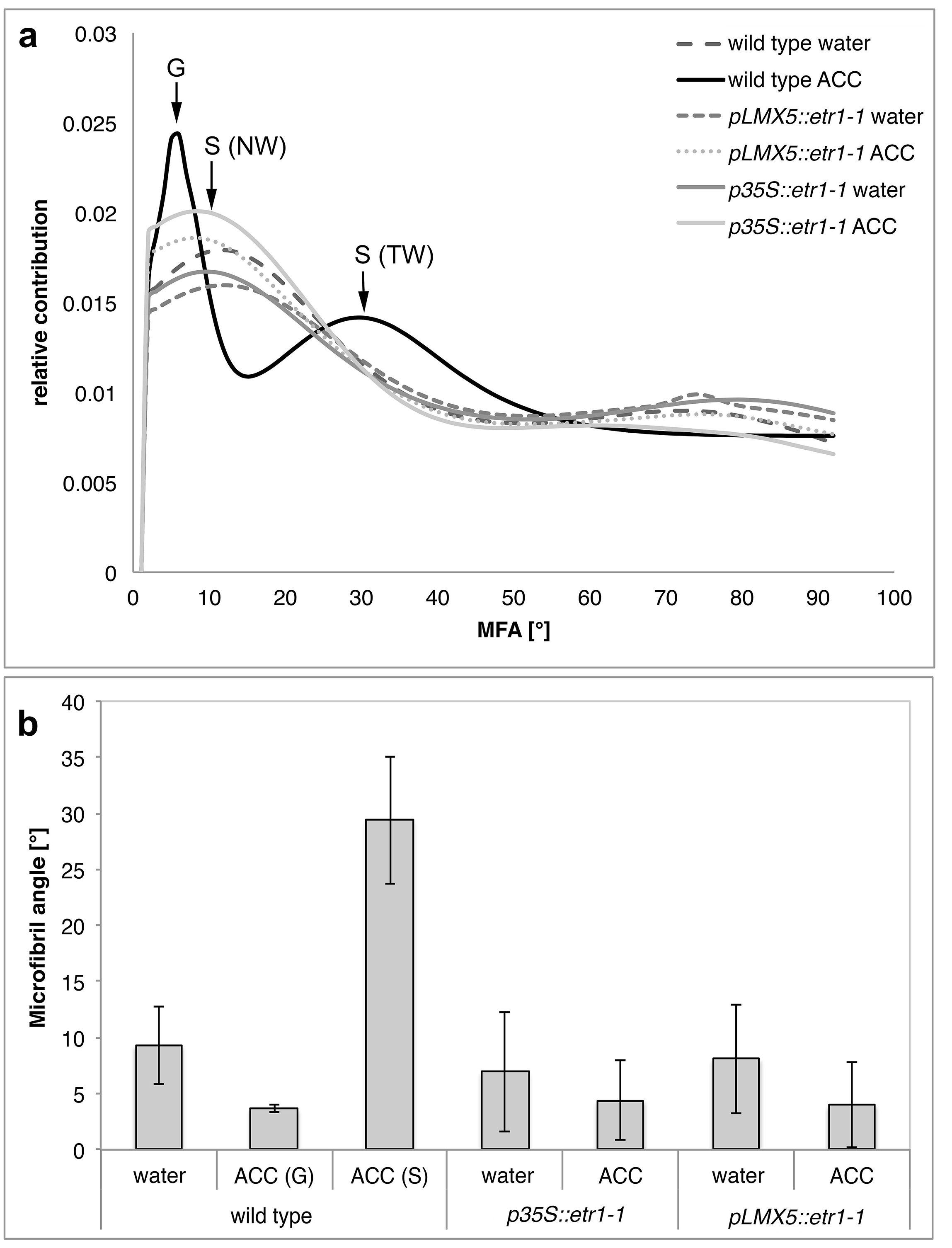
Cellulose orientation in ACC-and water-treated *P. tremula x P. alba* stems. (a) Cellulose microfibril orientation distribution shown as the relative contribution of microfibril angles (MFAs) to the overall distribution (sum of the contributions of 91 angles is 1). The orientation distributions represent mean distribution from all replicates per condition and genotype (n=5-7 with 2 technical replicates). Maxima for G-layer, S-layer in tension wood (TW; S (TW)) and in normal wood (NW; S (NW)) are indicated. Note the bi-modular pattern specific for ACC-treated wild type. (b) Mean microfibril angle of the different lines and treatments. G=MFA in G-layer, S=MFA in S layer. Error bars indicate ±SD (n=5-7).

To confirm that not only ACC, but also ethylene, can induce G-layer formation, greenhouse-grown trees were exposed to ethylene gas using flow-through cuvettes attached to the stem as described in Love et al. (2009). A two-week ethylene treatment significantly enhanced (more than two-fold) stem radial growth (Fig. S7) and induced visible differences in xylem anatomy of wild type trees (Fig. 5). A zone of fibers with thick cell walls was induced 1-1.5 mm from the cambium (Fig. 5c, 5g). From this zone radially outwards, typical ethylene effects were visible such as decreased vessels occurrence, with only few discontinuous arrangements of G-fibers. However, inwards from this zone of thick-walled fibers, G-fibers were induced. No similar cell wall modifications were observed in either the untreated wild type or in ethylene-treated or untreated ethylene insensitive trees (Fig. 5 and Fig. S8 and S9). Similar to the ACC-treated plants, G-fibers were transiently induced in a distinct ring around the stem, whereas reduced vessel occurrence was observed throughout the xylem formed after treatment. Because of the larger sized trees, and therefore the larger xylem area, it was possible to investigate the effect of ethylene on fibers without G-layers using an X-ray beam with a smaller diameter. These measurements showed that ethylene treatments induced an increased MFA in the region where most fibers were devoid of any G-layer (referring to the position magnified in Fig. 5c) as compared to control treatments, and that such effects were drastically reduced or absent in the ethylene insensitive trees (Fig. 6).

**Figure 5.**
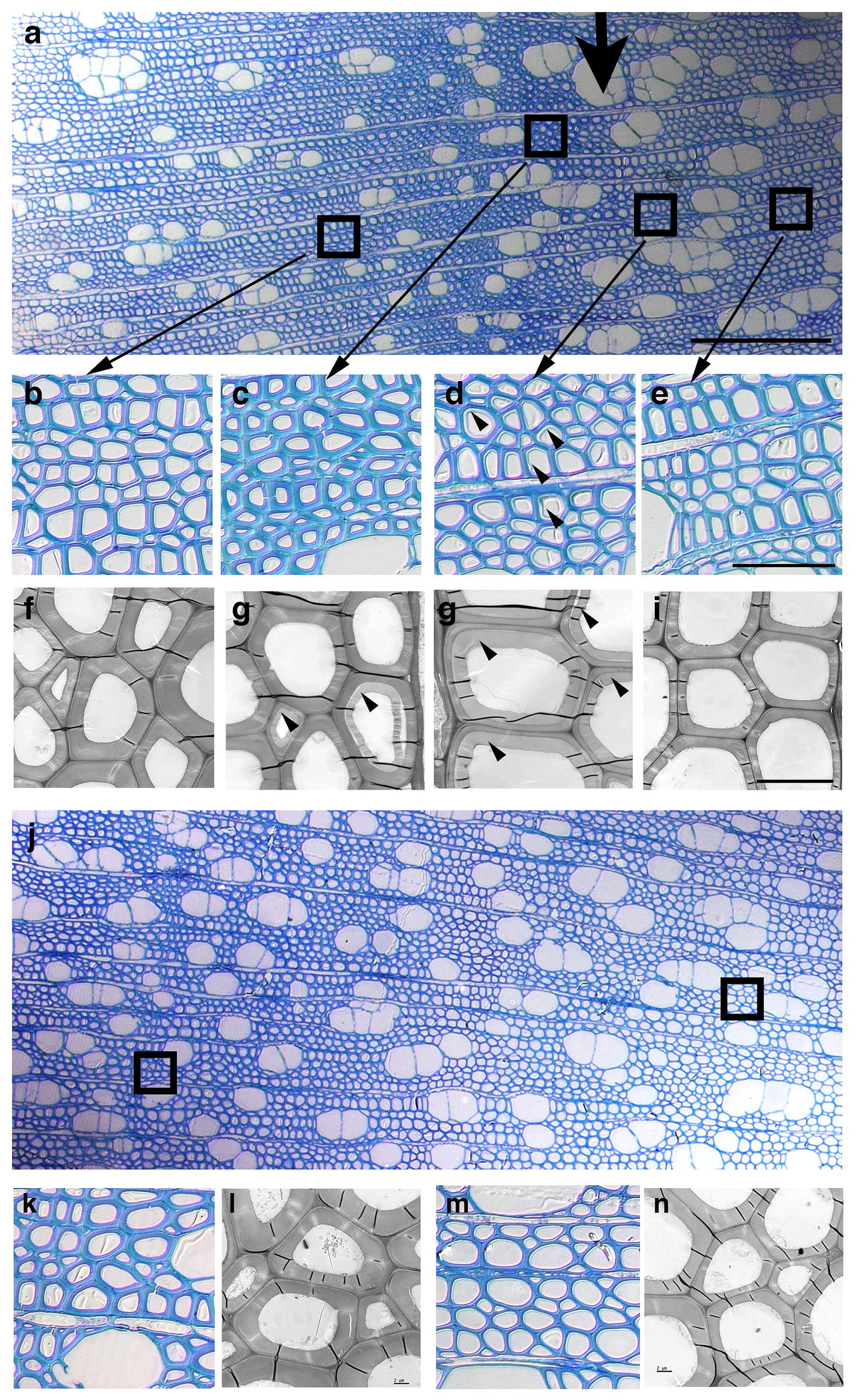
Light microscopy and TEM images showing gradients of cell wall thickness and patterning in greenhouse-grown *P. tremula x P. alba* wild type trees treated with synthetic air with ethylene (a-i) or with synthetic air without ethylene (control; j-n) by flow-through cuvettes. (a) Sections from the middle fragment of the ethylene-treated (a) and control (j) internode. Boxes in (a) and (j) show the positions that are magnified in the small images underneath each overview (b-i). Arrowheads in (d), (g) and (h) mark G-layers (in (d) only examples of cells with G-layers are indicated). Arrow in (a) indicates the boundary between xylem developed before (towards the right) and after (towards the left) ET treatment. Cambium is towards the left (a, j). Two positions, one more towards the cambium (left box in (j), magnified in (k), (l)) or towards the pith (right box in (j), magnified in (m), (n)) were selected to represent cell wall thickness in control conditions relative to ethylene-treated internodes. Scale bar in (a) and (j) is 200 μm, in (e) (also valid for b, c, d, k, m) is 50 μm and in (i) (also valid for f, g, h, l, n) is 10 μm.

**Figure 6.**
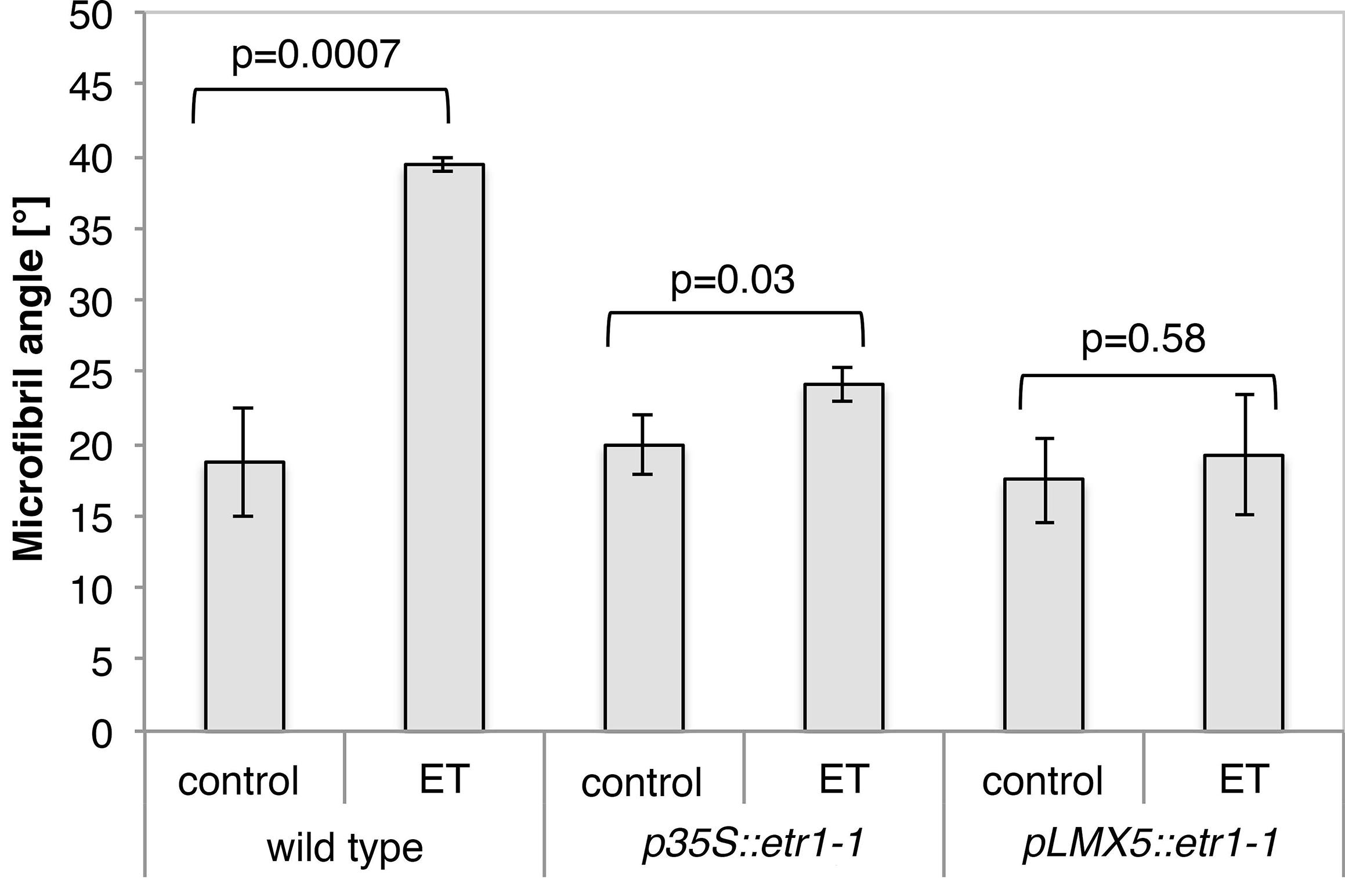
Average microfibril angle in greenhouse-grown *P. tremula x P. alba* wild type trees treated with synthetic air without ethylene (control) or with synthetic air with ethylene (ET) by flow-through cuvettes. Data were recorded within the first 400-500 μm inwards from the cambium. Error bars indicate ±SD (n=3). P-values of Student’s t-test between control and ethylene treatment conditions in the respective genotypes are indicated.

Taken together, exogenous ACC and ethylene induce the formation of G-fibers with G-and S-layers having MFAs typical for TW. TW characteristics such as decreased vessel occurrence and increased MFA in S-layers can occur in the absence of G-layers.

### Identification of genes downstream of ethylene signaling in stem tissues

We carried out an RNA sequencing analysis of stem tissues (comprising cortex, phloem, cambium, xylem and pith) of wild type and ethylene-insensitive trees treated with ACC or water for 10 h to identify potential targets of ethylene signaling for xylem development. The transcript profile in ACC-treated wild type plants was markedly different from all other samples (water treated wild type and ACC or water treated *p35S::etr1-1* and *pLMX5::etr1-1*) (Fig. 7 and Fig. S2 and S3). Only minor transcriptome changes were observed between ACC and water-treated trees in each ethylene-insensitive line, which again is in line with our previous findings that these lines are strongly but not totally ethylene insensitive. ACC induced a difference in transcript abundance of 1407 genes in wild type stems (≥ two-fold changes, padj≤0.01) (Table S2), 105 genes in *p35S::etr1-1* stems (Table S3) and 60 genes in *pLMX5::etr1-1* stems (Table S4). 1316 transcripts of the ACC induced genes (Table S5) were not significantly induced in the ethylene-insensitive lines (Fig. 7), and are therefore considered in the following as target genes of ethylene signaling. Transcripts of 23 genes (Table S6) showed significantly altered abundances in both wild type and ethylene-insensitive lines in response to ACC. The majority of these genes were, however, less induced in the ethylene-insensitive lines compared to the wild type. Therefore these transcripts could still be target genes in ethylene signaling considering the possibility that the ethylene-insensitive trees are probably slightly leaky.

**Figure 7.**
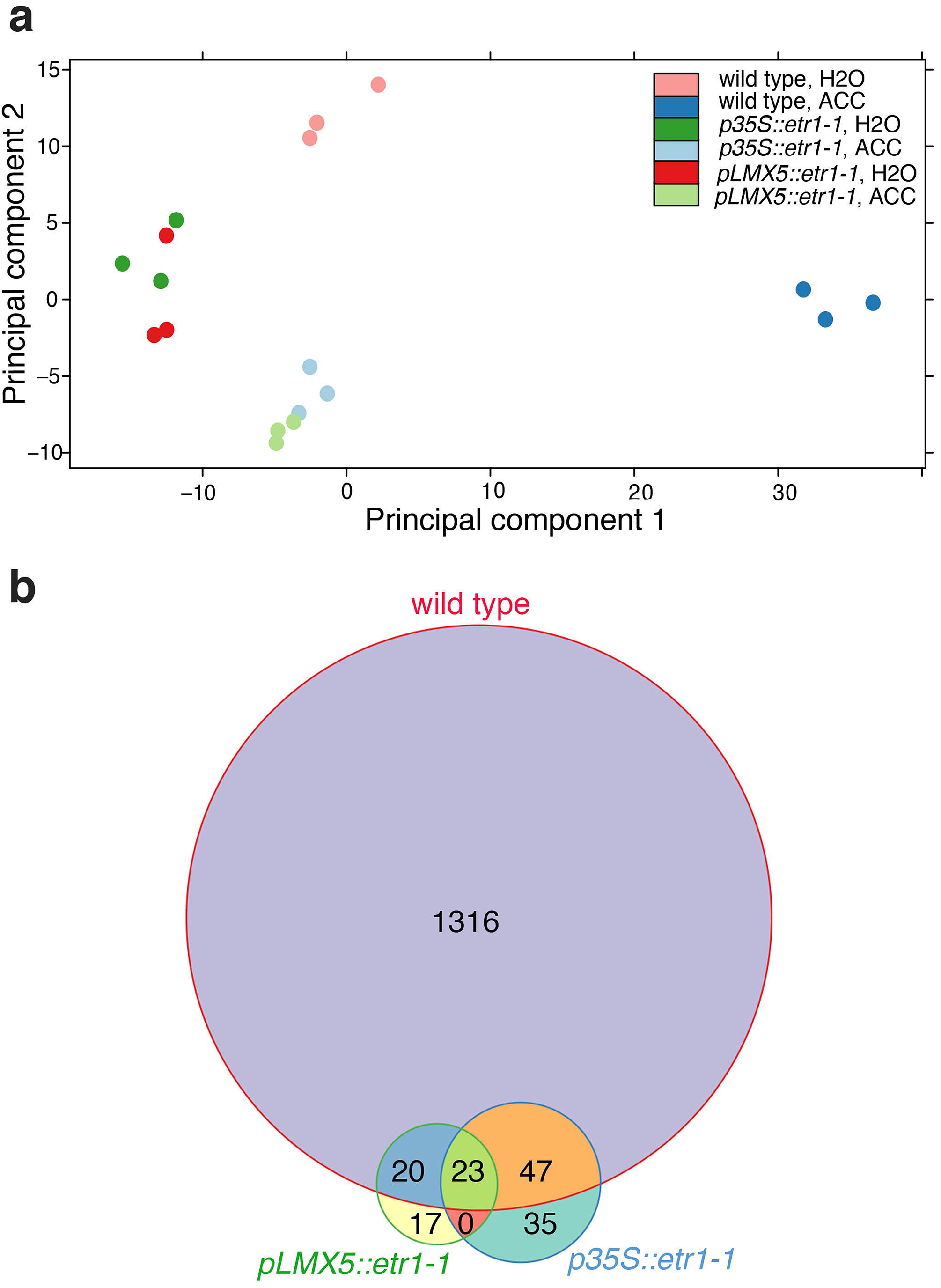
(a) PCA showing sample distances based on transcriptome analysis of different ACC-treated *P. tremula x P. alba* genotypes with respective water controls. (b) Venn Diagram of ACC-regulated genes with criteria padj ≤0.01 and fold change ≥2 in the respective genotypes after 10-h treatment compared to corresponding treatments.

It has been hypothesized that ACC has a signaling function independent of its signaling subsequent to conversion into ethylene (Xu *et al.*, 2008; Tsang *et al.*, 2011; Yoon & Kieber, 2013). Even though no obvious orthologs of the previously published ACC-responsive genes were detected in our data, three genes showed similar levels of induction in both ethylene-insensitive lines and wild type trees (<20% difference in fold change between the lines and wild type). These genes consist of two orthologs of the GA-responsive protein AtGASA1 (Potri.002G022600; Potri.002G022700), and an ortholog of Arabidopsis calcium-dependent protein kinase 24 (Potri.007G127000) (Table S6), which could be a potential regulator of ACC-synthase stability (Chae & Kieber, 2005; McClellan & Chang, 2008).

Genes with TF activity, among other categories, were overrepresented within the 1316 target genes according to a Gene Ontology (GO) enrichment analysis (Fig. S10). With help of the TF annotation in the plant TF database (PlantTFDB, http://planttfdb.cbi.pku.edu.cn/index.php?sp=Pth) we identified within our 1316 targets 168 TFs (Table S7), representing 12.7% of all 1316 target gene loci. For comparison, 6% of all gene loci in the *Populus* genome encode TFs, according to the number of genes annotated in PlantTFDB. Transcripts of 109 TFs were increased and 59 were decreased upon ACC treatment (Fig. 8). The largest category of increased transcripts (22 TFs) was the ERFs. Among these, 21 were also found to be induced two fold or more in response to ethylene treatment in hybrid aspen stems (Vahala *et al.*, 2013), evoking that these ERFs are indeed functional in ethylene signaling in *Populus* stems. Only four *ERFs* showed decreased transcript abundances, which is in agreement with the notion that *ERF* expression is positively rather than negatively influenced by ethylene signaling (Solano *et al.*, 1998; Feng *et al.*, 2005). In addition to the *ERFs*, a high number of *bHLH* (12 upregulated and 4 downregulated, 7.8% of all *bHLH TF* loci in the *Populus* genome), *MYB* (12 upregulated, 6 downregulated, 8.5 % of all *MYB TF* loci in the *Populus* genome), *NAC* (9 upregulated, 3 downregulated, 7.1 % of all *NAC TF* loci in the *Populus* genome) and *WRKY* (4 upregulated, 4 downregulated, 7.8 % of all *WRKY TF* loci in the *Populus* genome) (Fig. S11) TFs were present among our 1316 target genes. Eleven *bZIP TFs* were revealed as being differentially expressed but interestingly the majority of those transcripts were repressed upon ACC treatment (4 upregulated, 7 downregulated, 10.2% of all TF loci in the *Populus* genome (Fig. S11)). Lastly, three out of the seven loci that code for EIN3-like TFs in the *Populus* genome showed transcriptional induction upon ACC treatment. This is an unexpected result, as Arabidopsis EIN3 and EIN3-like are rather regulated on a posttranscriptional level (An *et al.*, 2010).

**Figure 8.**
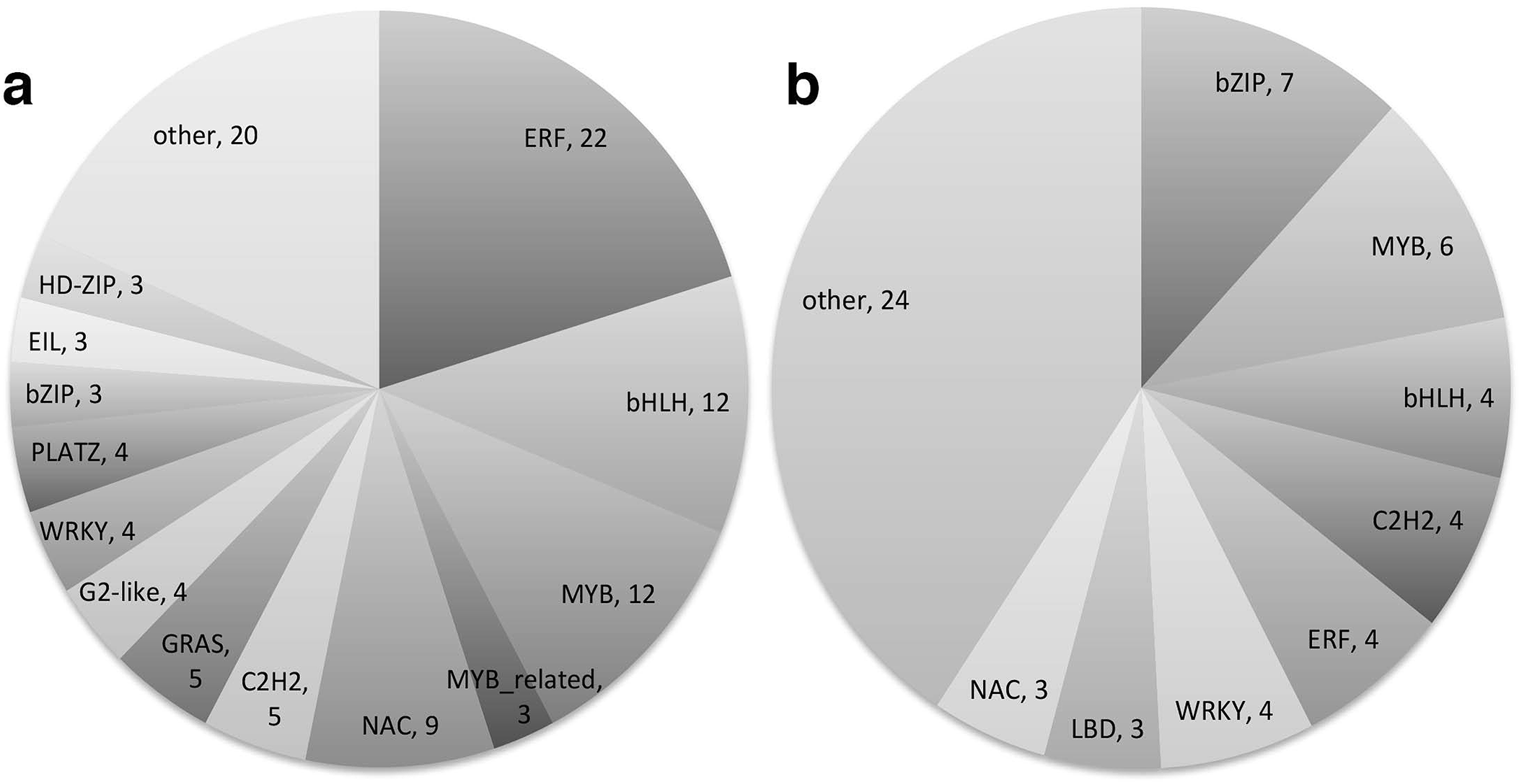
Classes of ACC-regulated transcription factors within the 1316 differentially expressed genes in *P. tremula x P. alba* wild type plants. Of the ACC-regulated transcription factors, (a) 109 were upregulated and (b) 59 were downregulated. Classification is based on Plant Transcription Factor Database, except for PLATZ transcription factors which were not in the classification. Numbers of ACC-regulated transcription factors are indicated. Those that were represented by only one or two members within their class are combined into the class “others”.

In order to assess crosstalk of ethylene with other hormones during G-layer induction, we screened the 1316 ethylene signaling target genes for hormone related genes (beyond the hormone-related TFs), using a keyword search within the description of their Arabidopsis orthologs. This identified 38 genes with increased expression and 25 genes with decreased expression (Table S9). The majority of these 63 genes were associated with auxin (23 genes), gibberellin (10 genes), ethylene (8 genes) and cytokinin pathways (6 genes). We also included in this list four genes related to polyamines that also function as plant growth regulators (Takahashi & Kakehi, 2010). The data show that the ethylene response in *Populus* stems is well established after 10 h of ACC-treatment: ethylene biosynthesis (ACC oxidase) is increased and the genes associated with the negative-feedback loop of the ethylene signaling pathway are induced as expected (Chang, KN *et al.*, 2013), e.g. homologs of *CONSTITUTIVE TRIPLE RESPONSE 1 (CTR1), ETHYLENE RESPONSE SENSOR 1 (ERS1), REVERSION-TO-ETHYLENE SENSITIVITY 1 (RTE1)* and *EIN3 BINDING F-BOX PROTEIN 1/2 (EBF1/2*). ACC treatment furthermore affected auxin related genes on all levels of auxin biology (biosynthesis, transport, conjugation and signaling) in woody tissues, and gibberellin-related genes regulating gibberellin catabolism, anabolism and signaling.

Finally, the expression of many genes related to cambial growth and cell wall formation was influenced by ACC treatments (Table S10). Three *PtCLE* genes *(PtCLE2, 27, 27)*, of which two, have, according to their expression, been associated with xylem tissues (Liu *et al.*, 2016), were found to be differentially regulated by ACC. Cytoskeleton related genes such as two AtMAP70-5 orthologs that are members of the actin and tubulin gene families as well as microtubule interacting proteins (TPX2 proteins) (Brunet *et al.*, 2004; Pesquet *et al.*, 2010; Goshima, 2011) were also affected. We observed slight induction of one gene associated with the Fascilin-like arabinogalactan (FLA) family, whose members have been reported to be highly induced during mature TW (Andersson-Gunnerås *et al.*, 2006) and sixty-three genes of the Carbohydrate Active Enzyme (CAZyme) family (based on published gene-lists by Geisler-Lee et al. (2006) (Table S10) related to cell wall biosynthesis and modification, exhibited altered expression. The large group of cell wall-related genes comprised genes encoding pectin degrading enzymes, including polygalacturonases, pectin methyl esterases (PMEs) and their inhibitors (PMEIs), and polysaccharide lyases. Several genes encoding expansins, xylogucan transglycosylases (XETs) and cellulases that are probably involved in cell expansion also exhibited altered expression after ACC treatment. Some polysaccharide biosynthetic genes from families GT2 with decreased expression and GT8 with increased expression were also affected.

### In silico *screening for potential direct targets of the ethylene signaling pathway in wood development*

To uncover putative direct target genes of the ethylene signaling pathway in wood formation, an *in silico* screening for EIN3 and ERF target motifs (TEIL and GCC box, respectively) was performed by analyzing the 2kb promoter region of all 1316 ethylene signaling target genes. Promoter regions were available in the Phytozome repository for 1292 of the 1316 genes. Generally, we found GCC boxes and TEIL motifs to be significantly enriched in the promoters of upregulated genes, but not in the promoters of downregulated genes. Of the 719 upregulated genes with available promoter sequences, 238 had TEIL motifs and 54 had GCC boxes within their 2kb promoter (for significant enrichment depending on promoter-length, see Table 1). Of the 585 downregulated genes only 12 harbored GCC boxes in their promoter, whereas 175 contained TEIL motifs (Table 1). The number of genes with motifs within the first 500bp upstream of the ATG was higher than the number of genes harboring motifs further away from the start codon (Fig. S12). Six ACC-regulated TFs had a GCC-box within the first 2kb of the promoter, while 42 contained TEIL motifs. (Table 1, Fig. S12). Interestingly, 15 ERFs (13 induced and two repressed ones) were identified as harboring TEIL binding sites in their promoter, suggesting that they are directly regulated by EIN3/EIL1 TFs (Table S11). Furthermore, 19 ethylene-regulated CAZymes have a TEIL motif in their promoter, 3 CAZymes harbor a GCC box and 1 gene harbors both a TEIL motif and a GCC box within the 2kb upstream of the start codon. Among these 24 genes, there were several pectin modifying genes that were strongly downregulated by ACC, as well as alpha and beta expansins both upregulated and downregulated.

**Table 1.**
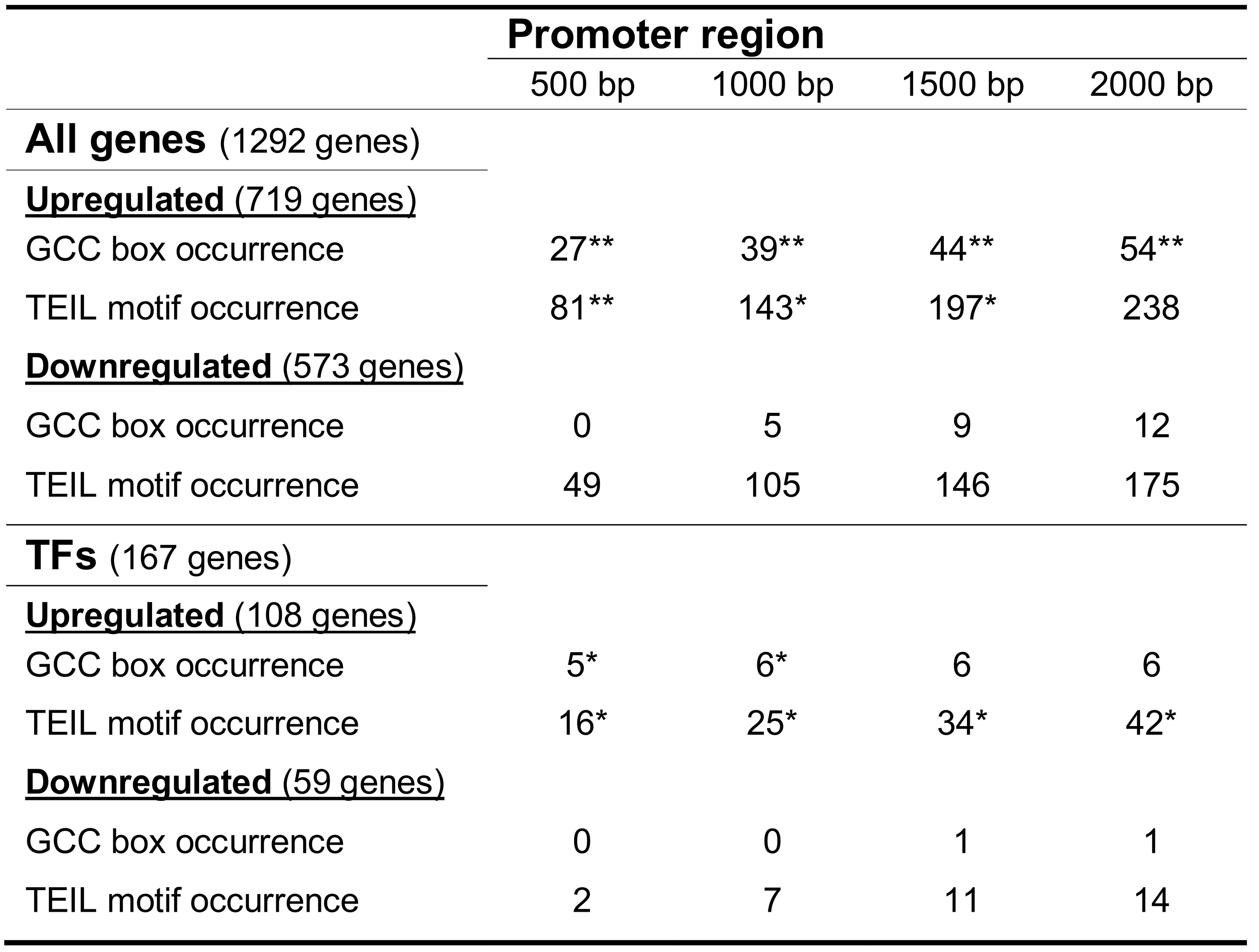
Number of genes with detected GCC or TEIL motifs in promoter regions of different lengths upstream of the start codon. Statistical enrichment compared to motif occurrence in promoters of all genes in the genome is indicated when applicable (*p-value < 0.05, **p-value < 0.01).

## Discussion

We demonstrated that ethylene signaling induces typical features of TW-G-fibers, including G-layers (Fig. 1 and 2), an altered chemical composition (Fig. 3) and a bimodal orientation distribution of the cellulose microfibrils (Fig. 4). Ethylene signaling also induced other features of TW such as enhanced cambial growth and decreased vessel size and frequency (Love *et al.*, 2009). This strengthens the concept that the increase in ethylene biosynthesis observed in association with stem displacement (Andersson-Gunnerås *et al.*, 2003) is a key component for triggering TW formation.

The induction of G-fibers in response to ethylene signals is most clear and consistent when ACC is applied through the xylem stream of *in vitro* grown plants. Possibly this experimental setup mimics best the situation in nature, and explains the lack of G-fiber formation in previous experiments using other methods of applying ethylene to the stem tissues (Yamamoto & Kozlowski, 1987; Junghans *et al.*, 2008). The exact site of ethylene production in woody stems is currently unknown. Experiments with *Zinnia elegans* cell cultures have suggested that ethylene production is an extracellular process (Pesquet & Tuominen, 2011). The ethylene precursor ACC is present in the xylem sap of aspen and its concentrations in the sap, as well as the abundance of ACC oxidase 1 activity in wood forming tissues, increases dramatically during TW formation (Andersson-Gunnerås *et al.*, 2003). This suggests a burst of ethylene during TW formation within the developing xylem.

G-fiber formation was transient during the experimental period of ACC- and ethylene-exposure (Fig. 1, Fig. 2 and Fig. 5), suggesting that the signaling/sensing conditions for induction of G-fibers changed as the experiment progressed. However the ethylene-stimulated decrease of vessel size and frequency was continuous throughout the treatment period. This indicates that G-fiber formation can be disconnected from other TW traits and that ethylene signaling can independently act on different stages of xylem cell development by interacting with the respective molecular machinery.

### Ethylene effect on microfibril orientation in secondary walls

ACC and ethylene applications led to an increase in the MFA in the S-layers of both G-fibers and non-G-fibers (Fig. 4 and Fig. 6). In primary cell walls, ethylene signals lead to a reorientation of cortical microtubules from transverse to longitudinal resulting in longitudinal microfibrils (lower MFA) (Apelbaum & Burg, 1971; Eisinger & Burg, 1972; Steen & Chadwick, 1981; Lang *et al.*, 1982). However, such effects have not been reported for the SCW. An increase in the S2-layer MFA of G-fibers has been reported previously (Goswami *et al.*, 2008; Clair *et al.*, 2011; Rüggeberg *et al.*, 2013). Even if the present study is not conclusive in determining exactly which S-layer the higher MFA in the non-G-fibers originates from, it can be concluded that ethylene has the capacity to alter the MFA of the secondary cell wall and that this process leads to opposite results (higher MFA) compared to the primary wall (lower MFA).

### Molecular links between ethylene signaling and wood formation

Transcription of numerous TFs is altered in response to ACC, with the largest proportion being ERFs (Fig. 8). ERFs present a possible molecular connection between the ethylene signaling pathway and wood development (Vahala *et al.*, 2013). Among the five ERFs that induced differences in wood chemistry when overexpressed (Vahala *et al.*, 2013), three (ERF18, ERF 21 and ERF85) were found to be induced by ACC in the present study. Of these, ERF18 and ERF21 harbored EIN3/EIL1 binding sites (TEIL motifs) suggesting a direct link to the ethylene-signaling pathway. Interestingly, the ERF18 and ERF21 over-expressing hybrid aspen trees were reported to contain elevated levels of carbohydrates and glycosidic linkages and reduced levels of lignin (Vahala *et al.*, 2013), which is in line with the now observed carbohydrate-rich G-fiber induction through ethylene signaling (Fig. 3). ERFs are therefore potential actors for the molecular connection of ethylene signaling to cell wall formation. Interestingly however, EIN3/EIL1 binding sites were more common than ERF binding sites (Table 1) in the promoters of all ACC-regulated, ethylene-downstream genes. This opposes the idea that ERFs (170 genes in *Populus*) downstream of EIN3/EIL1 (seven genes in *Populus*) would make the linear ethylene signaling pathway branch out in a vast spectrum of tissue- and process-specific responses. Both our data and a study by Chang et al. (2013), which aimed to identify targets of EIN3 in Arabidopsis via Chromatin-Immunoprecipitation, suggest that EIN3/EIL1 mediate a large panel of ethylene downstream responses independently of ERF signaling. Therefore targets of both ERFs and EIN3/EIL1 in wood development need to be considered to identify the molecular connection between ethylene signaling and wood formation.

### Regulation of cambial activity by ethylene

Ethylene application leads to enhanced cambial activity. The plant hormone auxin regulates cambial activity in trees in a dose-dependent manner (Sundberg and Little, 1990, Björklund et al., 2007). Auxin levels peak in the cambial zone in the wood, where cell division occurs, and decay rapidly towards the xylem and phloem side (Uggla *et al.*, 1996; Tuominen *et al.*, 1997; Uggla *et al.*, 1998). In response to ACC, 28 genes related to auxin metabolism, transport and signaling (incl. TFs) were differentially expressed (Table S9). These included three Small Auxin Up RNAs (SAURs), one potential auxin-conjugation enzyme (GH3), orthologs to the auxin transporter AtPGP1 and five auxin-related TFs (three ARFs and two Aux/IAA orthologs). The fact that auxin signaling (the majority of SAURs) as well as auxin biosynthesis (an anthranilate synthase ortholog) were induced upon ACC treatment suggests an increase in auxin during this early response to ACC, which is in line with the observation in Arabidopsis roots, that ethylene upregulates the transcription of auxin biosynthesis genes (Stepanova *et al.*, 2005; Stepanova *et al.*, 2008). However, none of the ethylene-regulated auxin genes carried a TEIL motif or GCC box in their 2kb promoter, defeating a direct molecular connection between ethylene signaling and auxin biosynthesis/signaling under our experimental conditions.

### Regulation of cell differentiation (fiber-to-vessel ratio) by ethylene

Interestingly, the majority of cell wall related genes that were affected by ACC were repressed (Table S7 and S10). The fact that many of these contained TEIL motifs in their promoters, proposes that they are directly suppressed by ethylene signaling. Despite EIN3/EIL1 usually being reported as an activator of gene expression, this TF has been shown to be able to function as a repressor, although the mechanism is not well understood (Shi *et al.*, 2012). Repressed genes may include vessel-specific cell wall biosynthetic/modifying enzymes, because the vessel cell differentiation is inhibited by ACC treatment. Cell wall related TFs have been characterized in *Populus* (Zhong *et al.*, 2010; Zhong *et al.*, 2011; Chai *et al.*, 2014; Wang *et al.*, 2014). Among these, the NAC TF *PtrWND2A*, homolog of *AtNST1* acting as a master regulator of secondary cell wall formation, was repressed upon ACC treatment. *PtWND2A* induces ectopic secondary cell wall formation when overexpressed in Arabidopsis or poplar plants (Ohtani *et al.*, 2011), although this was not observed in an earlier study (Zhong *et al.*, 2010). Furthermore, two MYB TFs with a potential connection to xylem formation were induced. *PtMYB192* is a close homolog to *AtMYB63* (Wang *et al.*, 2014), which regulates lignin genes in Arabidopsis (Zhou *et al.*, 2009). The poplar gene can be induced to a certain extent by the upstream secondary cell wall regulator *PtWND2B. PtMYB055* is highly expressed in stem tissues (phloem/cambium/xylem, (Chai *et al.*, 2014)), but as yet, no function has been reported for this MYB. Neither *PtWND2A* nor *PtMYB192* nor *PtMYB055* contain an ethylene motif in their 2kb promoter and are therefore unlikely to be regulated directly through the ethylene TFs.

### Ethylene induces changes in transcripts related to cell wall formation

Ten hour-ACC treatments altered the expression of many cell wall-related genes (Table S10). Many of these genes relate to cell expansion and cell wall modification. Pectin-related genes were mostly downregulated by ACC, for example the putative polygalacturonases (*GH28_6*, *GH28_46, GH28_73, GH28_75* and *GH28_79*), pectin methyl esterase (*CE8_67*), pectate lyases (*PL1_3, PL1_9* and *PL1_19*), and rhamnogalacturonan lyase (PL4_1), indicating decreased loosening of the pectin structure, which could explain reduced cell expansion. Similarly, genes encoding cell wall bound cellulases *PtCEL9B3* and *PtGH9B6* (Rudsander *et al.*, 2003; Takahashi *et al.*, 2009) and alpha-expansins, *PtEXPA5* and *PtEXPA15*(Gray-Mitsumune *et al.*, 2004; Sampedro *et al.*, 2006) were downregulated, whereas two other alpha- (*PtEXPA2* and *PtEXPA11*) and two beta- (*PtEXPLB3* and *PtEXPLB4*) expansins were upregulated. Among the *XTH* genes affected by ACC, one, *PtXTH30*, was downregulated, whereas two, others, *PtXTH34* and *PtXTH18*, were upregulated. These genes are thought to encode xyloglucan endotransglycosylase (XET) (Rudsander *et al.*, 2003; Baumann *et al.*, 2007). The products of these cell wall-remodeling related genes could be involved in cell wall modification in different stem tissues induced by ACC. A recent RNA sequencing study of fibers and vessel elements found that some of the downregulated genes (*PtGH28_73, PtGH28_79, PtCE8_67, PtPL1_3, PtPL1_19*, and *PtXTH30*) are highly expressed in developing vessel elements (Shi *et al.*, 2017). Thus, these genes could potentially be involved in the development of vessel element perforations and/or in vessel element cell wall modification related to cell wall expansion or lignification. Their downregulation by ACC could be a part of the inhibition of vessel element differentiation by ethylene (Love *et al.*, 2009). The transcriptome data revealed repression of two AtMAP70-5 orthologs, one of which harbored a TEIL motif in its promoter and could putatively be directly influenced by ethylene signaling. AtMAP70-5 has been reported to be involved in SCW patterning in tracheary elements and even xylem development and organization in Arabidopsis (Pesquet *et al.*, 2010), and could furthermore support the molecular origin of the altered fiber to vessel ratio during ethylene response. Upregulation of XETs by ACC is in line with the proposed involvement of XET activity in TW formation (Nishikubo *et al.*, 2007; Gerttula *et al.*, 2015) and the special role of xyloglucan in the mechanism of tension building in the tissue (Gorshkova *et al.*, 2015).

Taken together, many potential direct connections between ethylene-signaling and more downstream acting cell-wall genes were uncovered for both ERFs and EIN3/EIL1 transcription factors. Interestingly no direct connection between ethylene-signaling and key regulatory transcription factors of wood formation was evident. This reveals that ethylene may effect different aspects of wood formation in a more targeted way and can potentially bypass the molecular machinery of master regulators for xylem formation to alter wood development. EIN3/EIL1 TFs might participate in this direct regulation of target genes and not only influence wood development through ERF regulation.

## Accession numbers

Raw sequencing data are available at the European Nucleotide Archive (ENA) https://www.ebi.ac.uk/ena under the accession number ERP012528.

## Acknowledgements

J.F. and J.Lo. were funded by the Swedish Research Council Formas (Grant 213-2011-1148 and 239-2011-1915). We wish to acknowledge Gunilla Malmberg for preparation of plant material, Kjell Olofsson for help with sectioning, Lenore Johansson for sample preparation for transmission electron microscopy, Bastian Schiffthalter for help with DESeq transcript data analysis and David Sundell for help with generation of GO-enrichment plots on PopGenIE. The support of the Chemical Biological Centre and the Department of Chemistry, Umeå University, Sweden, for the Vibrational Spectroscopy Core Facility is gratefully acknowledged.

## Author contributions

J.F. carried out ACC treatment experiments and prepared material for analyses, carried out analyses, analyzed and interpreted the data and wrote the manuscript. J.V. carried out ethylene treatment experiments and contributed to writing the manuscript. J.Lo. carried out preliminary studies relating to the experiments presented. A.G. carried out FT-IR microspectroscopic measurements, data analysis and interpretation and contributed to writing the manuscript. M.R. carried out X-ray diffraction measurements, analyzed and interpreted the data and contributed to writing the manuscript. N.D. assisted with the bioinformatics analysis and contributed to writing the manuscript. J.Le. carried out immunolocalization experiments. J.K. contributed to data interpretation and writing the manuscript. T.R.H. carried out motif analysis and contributed to writing the manuscript. E.M. interpreted data from immunolocalization experiments, cell wall-related transcript changes and contributed to writing the manuscript. B.S. conceived and coordinated the study and wrote the manuscript.

## Supporting Material

### Methods

**Method S1**: Additional information on Plant Material, growth conditions and ACC treatment

**Method S2**: Immunolocalization procedures with LM5 and LM10 antibodies

**Method S3**: FT-IR microspectroscopy – technical details and information on multivariate data analysis

**Method S4**: Specifications on RNASeq data analysis

### Supporting Figures

**Fig. S1** Ethylene-effect in greenhouse-grown trees at three stem positions along the treatment area.

**Fig. S2** RNA-Seq data: Sample to sample distances of the respective replicates based on normalized read counts (DESeq) from RNA-Seq data.

**Fig. S3** RNA-Seq data: Log2 fold change against mean normalized counts and histograms of p-Value distribution.

**Fig. S4** Sections of biological replicates of water- and ACC-treated wild type and ethylene insensitive trees.

**Fig. S5** Successive TEM images captured in radial orientation from cambium to pith through cross-sections of wild type hybrid aspen xylem treated with water or ACC.

**Fig. S6** Immuno-gold localization of xylan (LM10 antibody) and of β-1,4-galactan (LM5 antibody) in stem cross-sections of ACC-treated wild type trees.

**Fig. S7** Stem radial growth in ethylene-treated or control greenhouse-grown trees.

**Fig. S8** Sections of biological replicates of controls and ethylene-treated greenhouse grown wild type hybrid aspen trees.

**Fig. S9** Sections of biological replicates of ethylene-treated greenhouse grown ethylene-insensitive hybrid aspen trees compared to one wild type ethylene treated tree.

**Fig. S10** RNA-Seq data: Enrichment of GO classes by molecular function in the 1316 differentially expressed genes.

**Fig. S11**: RNA-Seq data: Number of ACC-regulated transcription factors compared to the total number of loci for each transcription factor family in the *Populus* genome.

**Fig. S12**: Distribution of GCC box and TEIL motifs within the promoters of ACC-regulated genes.

### Supporting Tables

**Table S1**: Metadata for raw and post-QC read counts and alignment rates in RNA-Seq data

**Table S2**: Differentially expressed genes in ACC-treated compared to water-treated wild type hybrid aspen stems

**Table S3**: Differentially expressed genes in ACC-treated compared to water-treated *p35S::etr1-1* stems

**Table S4**: Differentially expressed genes in ACC-treated compared to water-treated *pLMX5::etr1-1* stems

**Table S5**: Differentially expressed genes downstream of ethylene signaling

**Table S6**: 23 commonly regulated genes in all three genotypes

**Table S7**: Differentially expressed transcription factors within the 1316 target genes

**Table S8**: Comparison between the gene expression of all ERFs from RNA-seq data and qPCR data from Vahala et al., 2013

**Table S9**: Hormone-related genes within the 1316 differentially expressed target genes

**Table S10**: Secondary growth or cell-wall-related genes within the 1316 target genes

**Table S11**: TEIL and GCC-box motifs within the first 2kb promoter of the 1316 target gene

